## Supplemental Materials for "Multiple memories can be simultaneously reactivated during sleep as effectively as a single memory"

Supplementary Materials include:

1. Supplementary Figure 1

2. Supplementary Figure 2

3. Supplementary Figure 3

4. Supplementary Table 1

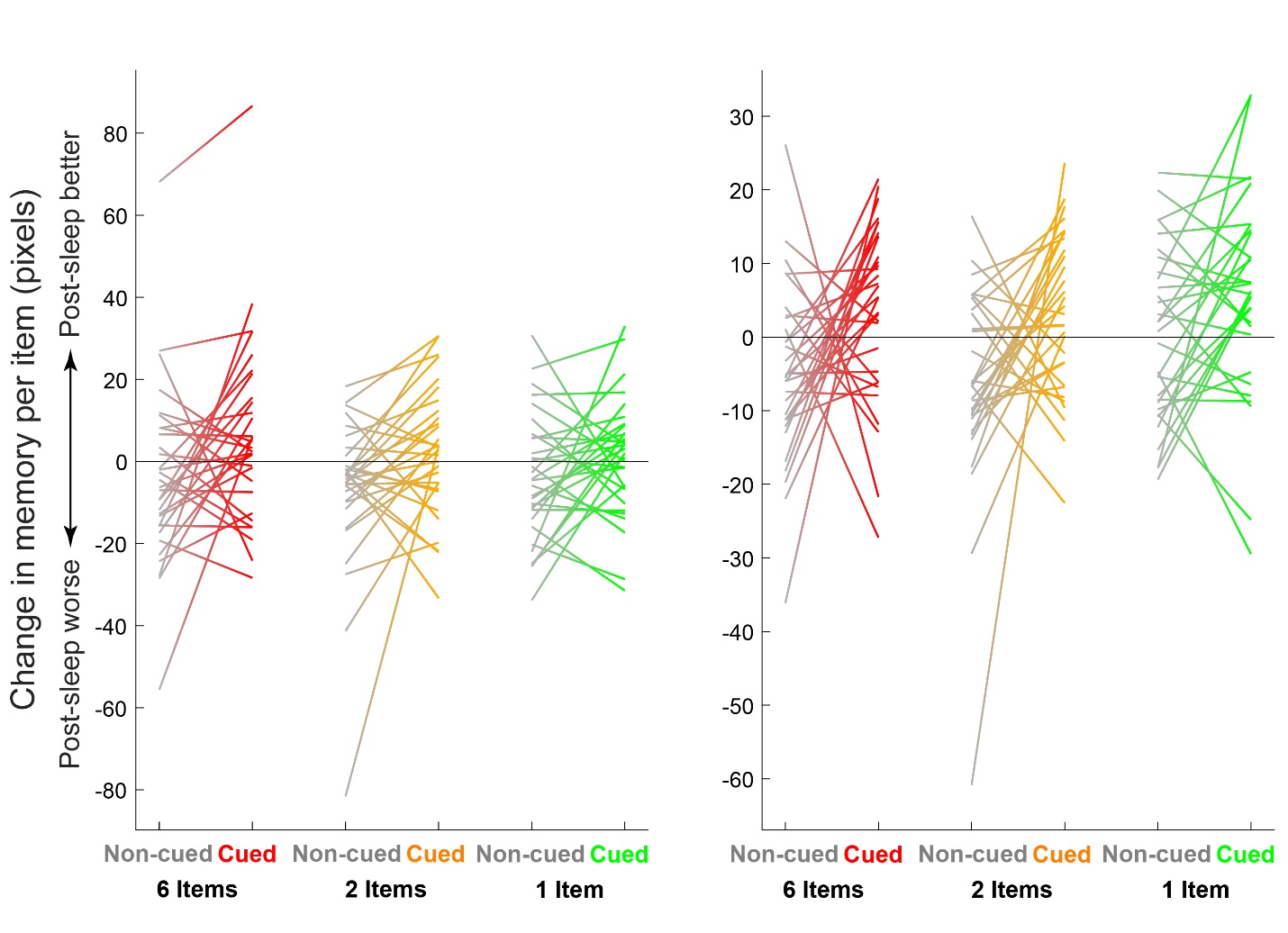

**Supplementary Figure 1: Individual participant averages of the change in accuracy over sleep for cued and non-cued items of different set sizes.** Results are shown for the uncorrected values used for the analysis shown in Figure 2 (left) and for the corrected values (in which the pre-sleep error rates were regressed out) used for supplementary analysis (right). Source data are provided as a Source Data file.

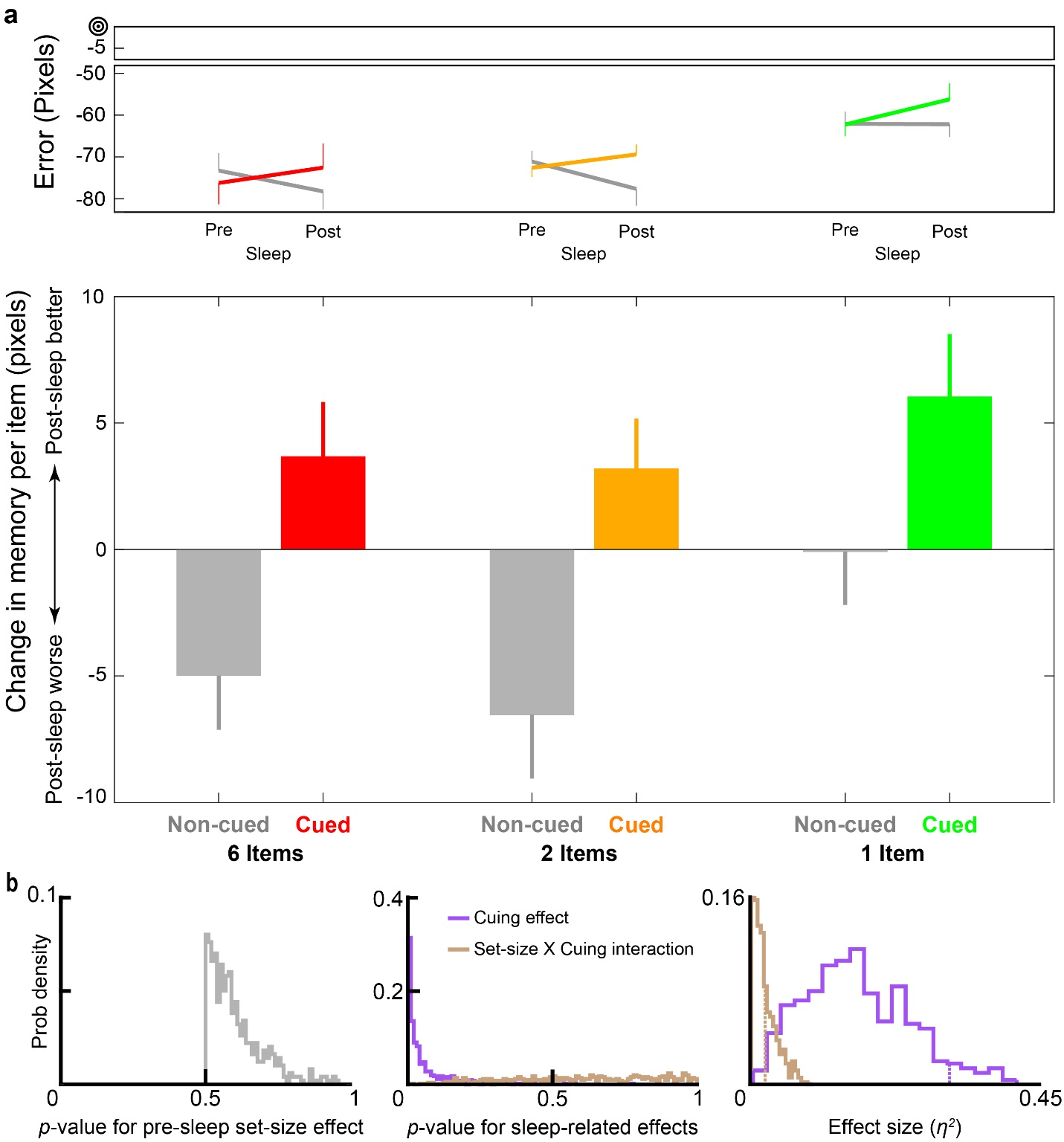
 **Supplementary Figure 2: The effect of cuing and the null interaction effect between cuing and set-size are not a result of pre-sleep differences in errors between set-sizes.** (a) To rule this option out, we regressed the pre-sleep error rates out. Note that pre-sleep values are therefore identical to those presented in Figure 2a. As for the uncorrected data, with the corrected data the cuing effect is significant (*p*<0.001) and the cuing-by-set-size interaction is not (*p*=0.75). (b) To complement this method, we subsampled data in a way that would minimize effects of pre-sleep differences (i.e., we chose a subsampled dataset that had a pre-sleep set-size difference with *p*>0.5, as shown in the histogram on the left). We then calculated *p*-values for the cuing effect and the cuing-set-size interaction for these datasets (histograms shown in center) and showed that the cuing effect was consistently significant whereas the interaction was consistently not. The right panel shows a histogram of the effect sizes (*η*^2^) of the cuing effect (purple) and the interaction effect (brown) for the subsampled data sets. Dashed vertical lines show the effect sizes calculated for the full datasets (i.e., without subsampling). Source data are provided as a Source Data file.

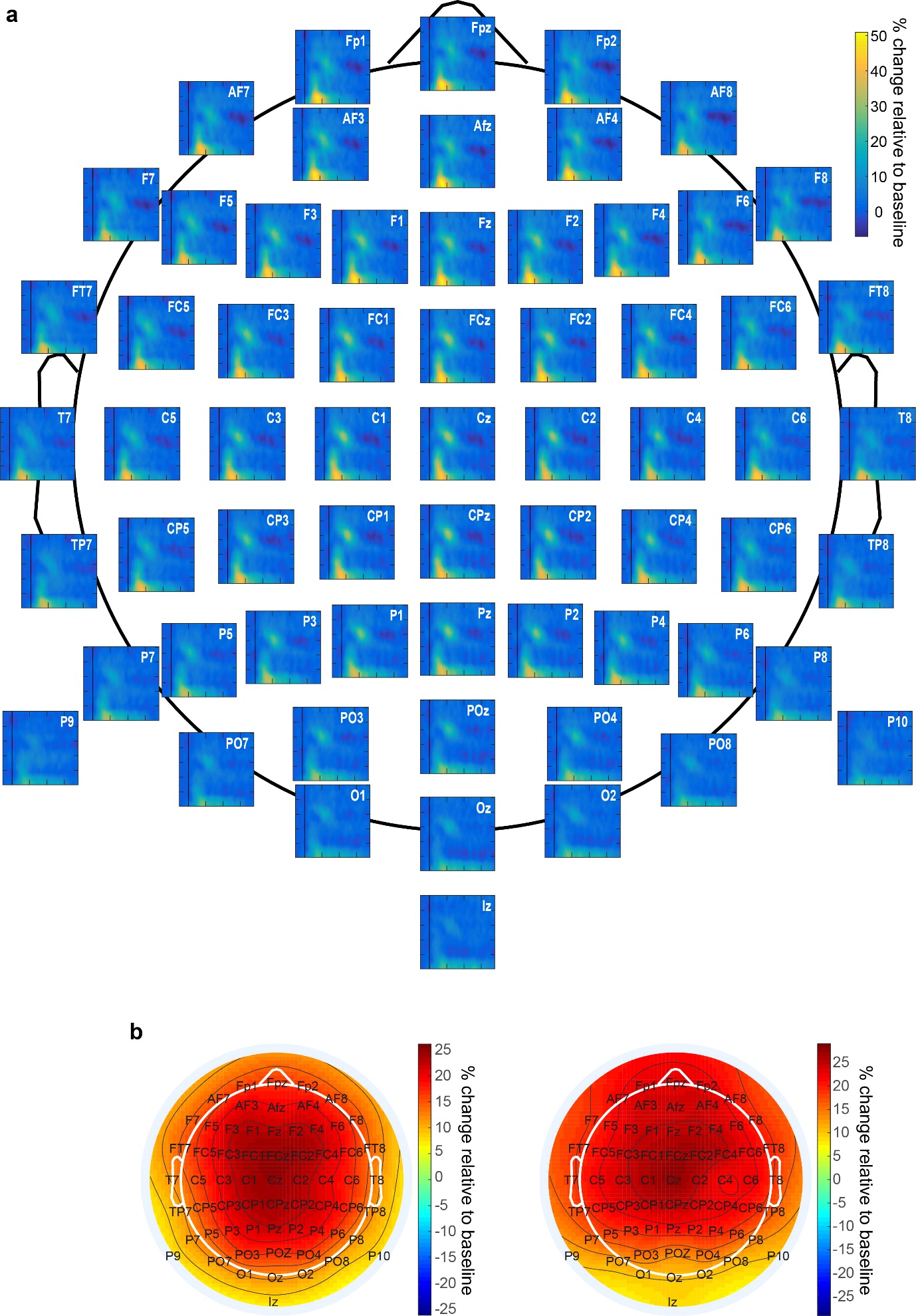

**Supplementary Figure 3: Additional physiological data.** (a) Spectrograms for all scalp electrodes, locked to cue onset during sleep and averaged over trials regardless of condition. Image positions on the scalp reflect electrode positions. (b) Scalp distributions of power modulations for delta-theta (left) and sigma (right), the two main clusters shown in Figure 4b. Source data are provided as a Source Data file.

**Supplementary Table 1: Themes used for sets of images**

| **Theme** | **Used as** |
| --- | --- |
| Balls | Two- or six-item sets |
| Fire | Two- or six-item sets |
| Birds | Two- or six-item sets |
| Documents and Literature | Two- or six-item sets |
| Cameras | Two- or six-item sets |
| Automobiles | Two- or six-item sets |
| Cats | Two- or six-item sets |
| Timepieces | Two- or six-item sets |
| Clothes with zippers | Two- or six-item sets |
| Dogs | Two- or six-item sets |
| Doors | Two- or six-item sets |
| Drinks | Two- or six-item sets |
| Food | Two- or six-item sets |
| Frogs | Two- or six-item sets |
| Heart | Two- or six-item sets |
| Kettles | Two- or six-item sets |
| Toilets | Two- or six-item sets |
| Trains | Two- or six-item sets |
| Phones | Two- or six-item sets |
| Musical Keyboard | Two- or six-item sets |
| Airplane | Two- or six-item sets |
| Cough | Two- or six-item sets |
| Flowers | Two- or six-item sets |
| Shoes | Two- or six-item sets |
| Pen | One-item sets |
| Trumpet | One-item sets |
| Violin | One-item sets |
| Monkeys | One-item sets |
| Kiss | One-item sets |
| Pen | One-item sets |
| Cow | One-item sets |
| Pig | One-item sets |
| Record | One-item sets |
| Money | One-item sets |
| Laugh | One-item sets |
| Toothbrush | One-item sets |
| Robot | One-item sets |
| Owl | One-item sets |
| Gong | One-item sets |
| Lobby bell | One-item sets |
| Drop | One-item sets |
| Boiling water | One-item sets |
| Slinky | Practice sets |
| Balloon | Practice sets |
| Computer keyboard | Practice sets |
